# On the dynamics of spatial updating

**DOI:** 10.1101/2021.10.27.465887

**Authors:** Jean Blouin, Jean-Philippe Pialasse, Laurence Mouchnino, Martin Simoneau

**Author notes:** CORRESPONDENCE: Jean Blouin.

## Abstract

Most of our knowledge on the human neural bases of spatial updating comes from fMRI studies in which recumbent participants moved in virtual environments. As a result, little is known about the dynamic of spatial updating during real body motion. Here, we exploited the high temporal resolution of electroencephalography (EEG) to investigate the dynamics of cortical activation in a spatial updating task where participants had to remember their initial orientation while they were passively rotated about their vertical axis in the dark. After the rotations, the participants pointed towards their initial orientation. We contrasted the EEG signals with those recorded in a control condition in which participants had no cognitive task to perform during body rotations. We found that the amplitude of the P_1_N_1_ complex of the rotation-evoked potential (RotEPs) (recorded over the vertex) was significantly greater in the Updating task. The analyses of the cortical current in the source space revealed that the main significant task-related cortical activities started during the N_1_P_2_ interval (136-303 ms after rotation onset). They were essentially localised in the temporal and frontal (supplementary motor complex, dorsolateral prefrontal cortex, anterior prefrontal cortex) regions. During this time-window, the right superior posterior parietal cortex (PPC) also showed significant task-related activities. The increased activation of the PPC became bilateral over the P_2_N_2_ component (303-470 ms after rotation onset). In this late interval, the cuneus and precuneus started to show significant task-related activities. Together, the present results are consistent with the general scheme that the first task-related cortical activities during spatial updating are related to the encoding of spatial goals and to the storing of spatial information in working memory. These activities would precede those involved in higher order processes also relevant for updating body orientation during rotations linked to the egocentric and visual representations of the environment.

## Introduction

The capacity to keep track of our position in the environment is paramount when moving around. This cognitive skill is generally referred to as spatial navigation. In humans, large advances on the neural bases of spatial navigation were obtained by measuring the cerebral blood flow, with fMRI scanners, of recumbent participants virtually moving in visual environments. Studies employing these techniques have revealed a consistent set of cortical activations during spatial navigation (Wolbers et al., 2008; Sherril et al., 2015, Nemmi et al., 2013; Balaguer et al., 2016; Vass & Epstein, 2017; Ekstrom et al., 2003, Hartley et al., 2003). Increased activations were found in areas responding to visual stimuli (striate and extrastriate visual areas), and in regions not strictly involved in visual processing yet having important higher-order functions for spatial navigation. These regions include the posterior parietal cortex, the temporal and frontal cortices which contribute, in varying degrees, to working memory, space perception and spatial representations.

The dynamics of the neural network underlying spatial navigation uncovered by fMRI studies is largely unknown. This is notably due to the hemodynamic response time (Ghuman & Martin, 2019) which is too slow with respect to the speed of the processes engaged during spatial navigation (e.g., < 1.5 s for simple spatial updating tasks, Boon et al., 2018; Hodgson & Waller, 2006; Rieser, 1989). One can reasonably expect that the dynamics are conditioned, to some extent, by the functions of the different elements comprising the engaged network. For instance, early activation could be found in the areas which contribute to maintaining spatial information in short-term working memory (e.g., dorsolateral prefrontal cortex (dlPFC), see Wager & Smith, 2003; Gilbert & Burgess, 2008) and to processing external spatial information, such as goal destination (e.g., anterior PFC (aPFC), see Ekstrom et al., 2003; Ciaramelli, 2008; Spiers, 2008; temporal cortex, see Ekstrom et al., 2003, Hartley et al., 2003). On the other hand, later activations are to be expected in regions involved in higher cognitive processes. This could be the case for regions that contribute to the building of frames of reference that allow individuals to either encode the environment relative to themselves (e.g.,precuneus, Byrne et al., 2007) or to encode their position relative to the environment (lateral occipital cortex (LOC), Committeri et al., 2004; Zaehle et al., 2007).

Electroencephalography (EEG), with its excellent temporal resolution and the possibility to increase its spatial resolution using source analyses techniques (Im et al., 2007; Tadel et al., 2011, 2019), appears well adapted to capture the time course of spatial navigation (Ertl et al., 2017; Gale et al., 2016; Gutteling et al., 2015, 2016; Schneider et al., 1996). Moreover, the use of EEG also enables the investigation of brain activity in moving participants, i.e. where vestibular inputs provide the brain with critical body motions information for spatial navigation (Brandt et al., 2005; Schöberl et al., 2021; Kremmyda et al., 2016, see Smith 2017, for a review).

Gutteling et al. (2015, 2016) recently used EEG to record cortical activities of participants who had to retain the location of a peripheral target during passive whole-body motion in the dark (see Medendorp & Selen 2017, for a review). Investigating these activities in such spatial updating task can be thought of as a valuable entry point for getting insight into the cortical implementation of spatial navigation. Performed relatively well in darkness (see Klier & Angelaki, 2008 and Medendorp, 2011, for reviews), even in extreme cases of somatosensory deafferentation (Blouin et al., 1995), such tasks involve vestibular information processing. Gutteling et al. (2015, 2016) found a large alpha power decrease in electrodes overlaying the posterior parietal cortex (PPC) during spatial updating. Interestingly, the decreased alpha power was always located in the contralateral hemisphere to the memorised target and switched hemisphere when the unseen target changed visual hemifield during body motion. Reflecting enhanced cortical excitability (Jasper & Penfield, 1949; Pfurtscheller & Da Silva, 1999), the decrease of alpha power observed during the actual body motion provided human electrophysiological evidence that the PPC is involved in spatial updating (Duhamel et al., 1992; Medendorp et al., 2003; Merriam et al., 2003; Ventre-Dominey & Vallée, 2007) and in directing attention to locations or objects in the environment (Corbetta & Shulman, 2002).

Apart from the PPC, no other region of the spatial navigation network revealed by human fMRI investigations (see above) showed task-related neural oscillations in Gutteling et al.’s (2015, 2016) studies. This could be due to the fact that the EEG spectral content was examined in the electrode space (scalp level) rather than in the sources space. By partially de-convolving the EEG data in a physically and anatomically meaningful way, source space analyses may indeed reveal effects that remain undetected at the scalp level with electrode space analyses (Baillet et al., 2001; Salmelin & Baillet, 2009).

The goal of the present study was twofold: to analyse the cortical network involved in spatial updating during actual whole-body motion in the dark, and to obtain insight into the dynamics of this network. We performed source analyses of the EEG activity recorded while human participants were maintaining their initial orientation in memory while being rotated in darkness. The dynamics of the early stage of spatial updating (i.e., before predominance of processes related to the use of the updated spatial representation) was assessed by computing the current amplitude over the cortical surface in 3 consecutive time windows. These were defined by the negative and positive deflection points of the rotation evoked potential (RotEPs) recorded over the vertex. We predicted that the first task-related activations should be observed in areas involved in the short-term spatial working memory (e.g., dlPFC; Wager & Smith, 2003) and the online spatial updating processes (e.g., PPC; Duhamel et al., 1992; Gutteling et al., 2015, 2016). Later activations should be observed in regions involved in the egocentric encoding of spatial positions (e.g., precuneus, Byrne et al., 2007).

## Materials and methods

The data were collected in a previous study (Blouin et al., 2019). In this study, we specifically investigated the cortical activation associated with the planning of pointing movements whose targets were defined by idiothetic information issued from body rotations in the dark. This activation was assessed from the end of the rotation to the onset of the pointing movements (i.e., movement planning process). In the present study, we investigated the dynamics of cortical activation during the actual body motion (i.e., spatial updating process).

## Participants

Ten healthy right-handed participants (3 women, mean age: 26.6 ±2.7 years) participated in the experiment. They all had normal or corrected-to-normal vision and did not report any history of neural disorders. The data of one male participant had to be discarded because of technical problems. The experiment was conducted in accordance with the Declaration of Helsinki (except for registration in a database) and was approved by the Laval University Biomedical Ethics Committee. Informed consent was obtained prior to the experiment.

## Experimental set-up

The participants were seated in a dark room with their feet resting on a footstool. They were secured to the chair with a four-point belt. The chair could be manually rotated about its vertical axis by the experimenter. The rotations were recorded with an optical encoder at 1 kHz. A circular array of LEDs fixed on the floor behind the chair indicated its initial angular position and the 3 targets rotations (i.e., 20°, 30° and 40° in the counterclockwise direction). A light emitted by a laser diode fixed on the back of the chair provided the experimenter with visual feedback about the chair orientation along the LEDs array. The use of different rotation amplitudes together with the variability in the actual body rotation (e.g., acceleration, amplitude) for a given rotation target amplitude increased the necessity for the participants to direct their attention on information related to self-rotation to keep track of their initial orientation. Similar setups have frequently been used for testing vestibular-related processes (e.g., Hanson & Goebel, 1998; Blouin et al., 1998, 2010; Funabiki & Naito, 2002; Mackrous et al., 2019). Importantly, the choice of manual rotations reduced the possibility of electric noise contamination of the EEG recordings (see Nolan et al., 2009 for a discussion on this issue).

## Experimental tasks

### 1. Updating

Before the start of each trial, participants positioned their right hand on their ipsilateral knee and gazed at a chair-fixed LED positioned ∼1 m in front of them. They were instructed to keep fixating this LED during the whole duration of the trials. The participants received the preparation signal “ready” 2-3 s before either the 20°, 30° or 40° floor LED lit up behind the chair to indicate, to the experimenter, the amplitude of the next rotation. Some 100 ms after the end of the rotation, a beep indicated to the participants to produce a rapid lateral arm movement to point towards their pre-rotation, initial, orientation. The signal from the chair optical encoder was used to detect online the end of the rotation which was defined as angular velocity smaller than 2.5 °/s. The buzzer emitting the beep was located directly above the participants, along their longitudinal axis. This prevented the sound from providing spatial information (e.g., initial orientation). The participants were returned to the starting position after their pointing response and the next trial started following a minimum resting time of 15 s. Results related to the motor performance have been published separately (Blouin et al., 2019). Briefly, the burst of the arm muscular activities triggering the pointing movements occurred ∼400 ms after the imperative signal (beep) and the amplitude of the movements was scaled according to the amplitude of the body rotations. These behavioral results indicate that spatial updating was fast and attuned to the actual body rotation in space.

### 2. Control

We performed a second experimental block of trials like those performed in the Updating task, but with the only distinction that the participants did not produce arm movement when hearing the beep after the rotation offset. For these trials, the instruction was simply to keep ocular fixation on the chair-fixed LED during the rotations. This block of trials was used to normalize EEG activities recorded in the Updating task. As the dynamics of the body rotations were similar in both the Updating and Control tasks, as it will be demonstrated below, this normalization (also detailed below) allowed EEG activities that are not strictly related to spatial updating to be to cancelled out (see Gutteling et al., 2015). These activities might result for instance from eye movements, and from somatosensory (see Ert & Boegle, 2019) and vestibular stimulation during body rotations.

Participants were submitted to 25 rotations for each angular target for a total of 75 trials in each task (i.e., Updating and Control). The order of rotation amplitude was pseudo-randomly selected and the presentation order of the tasks was counterbalanced across participants.

On average (Control and Updating tasks), the rotation amplitudes were 19.81 ±0.83°, 29.67 ±0.96° and 40.53 ±0.94° for the 20°, 30° and 40° rotation tasks, respectively. To verify if participants experienced similar idiothetic information between the Updating and Control tasks, we compared chair angular acceleration between both tasks. To make this test, we first normalised the time-series of angular acceleration from start (0 %) to end (100 %); using an angular velocity threshold of 2.5°/s to identify beginning and ending of rotation. Then, for each chair rotation amplitude (i.e., 20°, 30°, 40°), we performed a two-tailed paired t-test using statistical parametric mapping (SPM) analyses. SPM uses random field theory to adjust for multiple comparisons. It enables comparison of continuous variables at time points other than discrete local optima (Pataky et al., 2013). This statistical approach is suited to analysing time-series where each sample is dependent on previous data points, as for acceleration data. For each rotation amplitude, results of the statistical tests revealed that the time-series acceleration did not significantly differ between the Updating and Control tasks (all Ps < 0.05; Fig. 1). These results suggest that if the dynamics of cortical activities differed between the Updating and Control tasks, this difference more likely would result from a difference between the cortical processes engaged within the tasks than from different idiothetic information.

**Figure 1.**
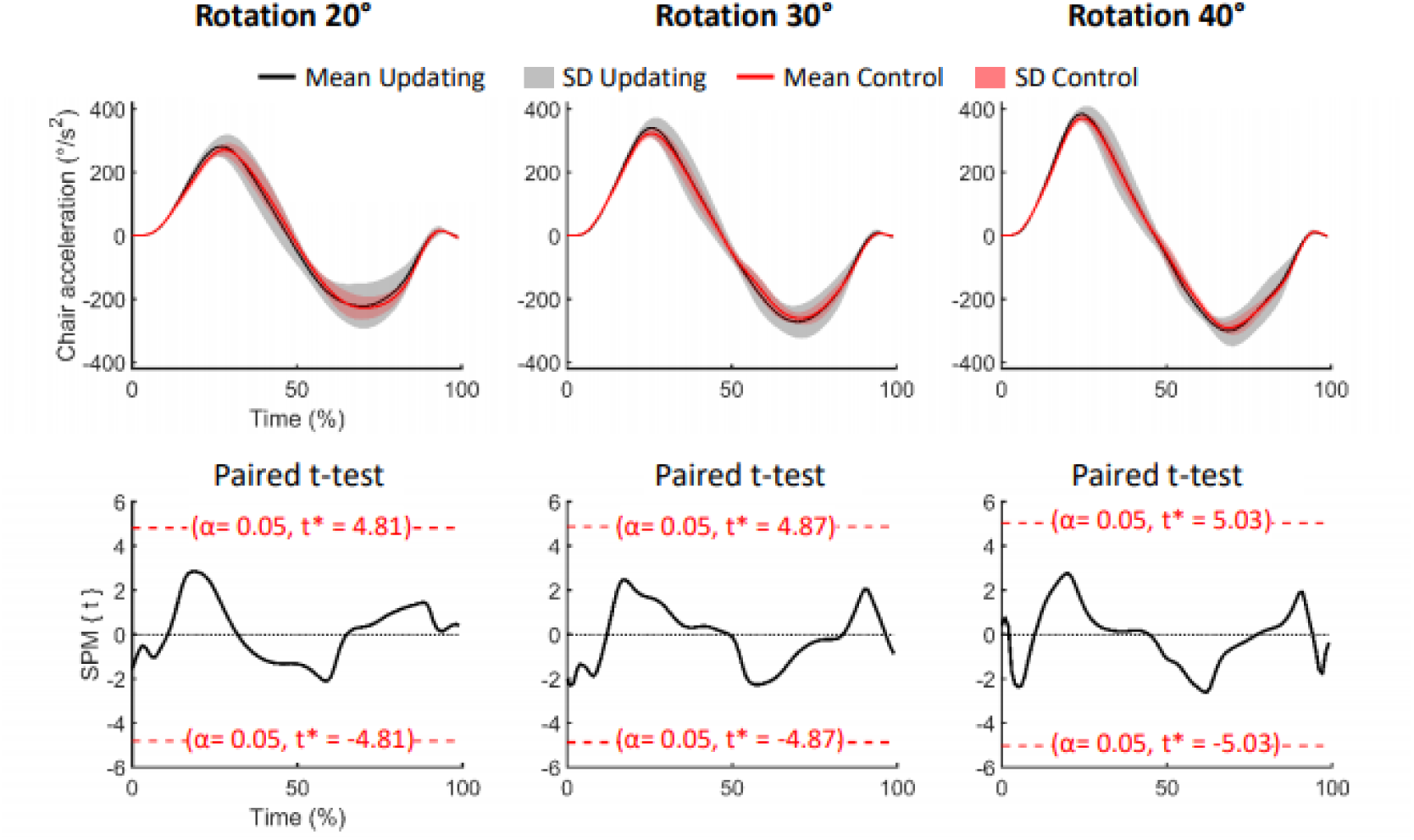
Data related to the chair rotations. Upper panels: mean (solid lines) and standard deviation (5D, areas) of the acceleration of the chair for the 20 ° (left), 30 ° (middle) and 40 ° (right) rotation amplitude. Black time-series depict the data for the Updating condition while the red time-series depict the data for the Control condition. Lower panels: results of the paired t-test for each rotational amplitude. SPM{t} represents the temporal trajectory of the t statistic (black lines) and the critical threshold (red lines, alpha = 0.05). T-score for each comparison is illustrated in each panel.

## EEG analyses

### Electroencephalographic (EEG) activity was recorded using a Geodesic 64-channel EEG sensor net (1000 Hz, Electrical Geodesics Inc., USA)

Data pre-processing was performed using BrainVision Analyzer 2 (Brain Products, Germany). The recordings were first referenced to the averaged activity of the 64 scalp electrodes. Then, data recorded by all electrodes were synchronized with respect to the onset of the rotation (i.e., when chair angular velocity > 2.5 °/s), with the average amplitude of the 200-ms pre-rotation epoch serving as baseline. Independent component analyses (ICA) were used to subtract ocular artifacts (e.g., blinks, saccades) from the EEG recordings. The recordings were visually inspected and epochs still presenting artifacts were rejected. The data were separately averaged for each participant, task, target body angular rotation amplitude (i.e., 20°, 30°, 40°) and electrode. These averages were used to estimate the sources of the cortical activities.

The cortical sources were reconstructed using Brainstorm software (Tadel et al., 2011, freely available at http://neuroimage.usc.edu/brainstorm). We employed the minimum-norm technique to resolve the inverse problem with unconstrained dipole orientations. The forward models were computed using a boundary element method (BEM, Gramfort et al., 2010) on the anatomical MRI Colin 27 high resolution brain template (306 716 vertices) provided by the Montreal Neurological Institute (MNI).

Consistent with previous studies (Schneider et al., 1996; Gale et al., 2016; Ertl et al., 2017), RotEPs were found over a large set of electrodes, but were largest over the Cz electrode (i.e. vertex). As shown in Fig. 2, the RotEPs were composed of successive inflection points which we refer to as P_1_ (over all mean 47 ±12 ms), N_1_ (136 ±12 ms), P_2_ (303 ±35 ms) and N_2_ (470 ±55 ms). These points served as temporal landmarks for analyzing the dynamic of the cortical activities in the source space.

**Figure 2.**
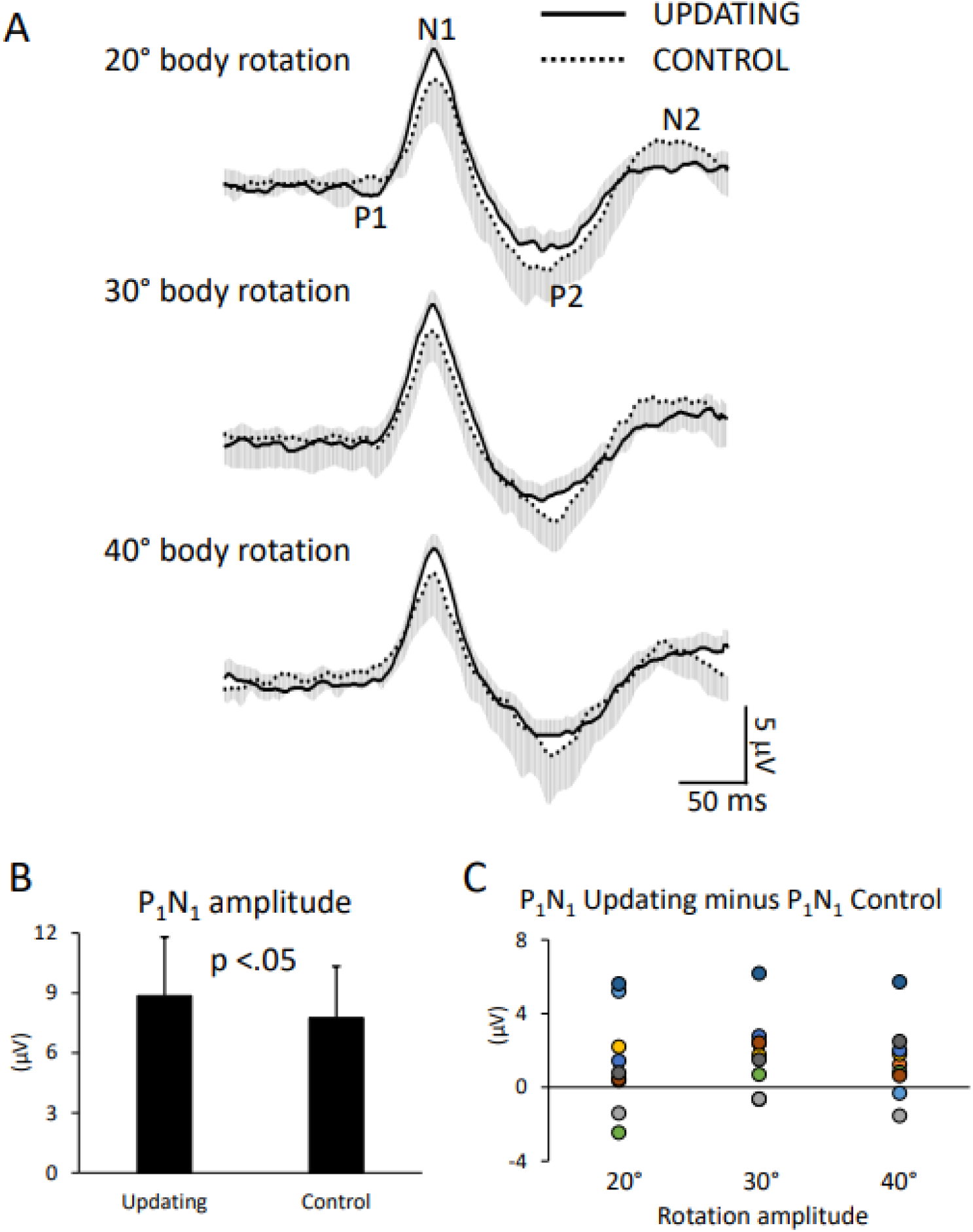
Data related to the rotation evoked potentials (RotEP). (a) Grand average RotEP traces at electrode Cz (i.e., vertex) in the Updating and Control conditions for the 20 °, 30 ° and 40 ° body rotations. Overall mean peak latencies: P_1_: 47 ±12 ms; N_1_: 136 ±12 ms; P_2_: 303 ±35; N_2_: 470 ±55 ms. (b) Mean P_1_N_1_ amplitudes in the Updating and Control conditions. Error bars represent the standard error of the means (c) Difference between the P_1_N_1_ amplitudes computed in the Updating and Control conditions. Dots of the same colour represent data from the same participant.

For each participant, rotation amplitude and task, we measured the latency of each RotEP’s inflection point and then averaged the current computed in the source space over 3 successive time windows: P_1_N_1_, N_1_P_2_ and P_2_N_2_. Following this step, we collapsed (by averaging) the maps obtained for each amplitude of rotations to obtain a single map of current amplitude per participant and task. It should be noted that this method for estimating EEG sources is not impacted by potential effects of rotation amplitudes, task or inter-individual differences on RotEPs peak latencies. Computed in the source space, current amplitude is considered to reflect brain activation (Tadel et al., 2011, 2019). Importantly, as the last RotEP peak considered in the Updating task (i.e., N_2_) occurred on average 171 ms, 269 ms and 326 ms before the end of the 20°, 30° and 40° rotations, different cortical activations between the Updating and Control tasks would unlikely stem from the mere preparation of the arm motor commands in the Updating task.

To highlight those cortical regions specifically involved in spatial updating, we computed statistical maps by submitting the current maps obtained in the Updating and Control tasks to t-tests (significance threshold p < 0.05). These analyses were performed separately for each time windows (i.e., P_1_N_1_: short latency, N_1_P_2_: mid-latency and P_1_N_2_: long latency) to gain insight into the spatio-temporal dynamics of the cortical activities during spatial updating. Sources were identified according to Brodmann’s areas after converting MNI to Talairach coordinates with the Nonlinear Yale MNI to Talairach Conversion Algorithm (Lacadie et al., 2008a). The Brodmann’s areas definition was based on Lacadie et al. (2008b).

The amplitude of the cortical potentials increases when they are evoked by task-relevant somatosensory (Cybulska-Klosowicz et al., 2011; Saradjian et al., 2013) or visual (Lebar et al., 2015) events. In the present study, extracting the RotEPs out of the EEG recordings allowed us to determine if this effect generalized to cortical responses evoked by idiothetic information relevant for spatial updating. To this end, we compared the amplitude of the P_1_N_1_, N_1_P_2_ and P_1_N_2_ components between the Updating and Control tasks. We also compared between these tasks, the latency of the RotEPs, which was defined as the time elapsed between the onset of the body rotation and P_1_. Variables related to the RotEP were submitted to separate 2 (Task: Updating, Control) by 3 (Amplitude: 20°, 30°, 40°) repeated-measures ANOVAs (significance threshold p < 0.05). Prior tests (Kolmogorov-Smirnov) confirmed the normality of all data.

## Results

### Rotation-evoked potentials (RotEPs)

The ANOVA revealed a significant effect of Task on P_1_N_1_ amplitude (Fig. 2). The amplitude of this first RotEP component was greater in the Updating than in the Control task (8.83 ±3.18 mV vs 7.76 ±3.14 mV, F_1,8_ = 5.66, p=0.04). The analyses did not reveal significant effect of Amplitude (p=0.38) or significant interaction of Task x Amplitude (p= 0.09). The other RotEP components (i.e., N_1_P_2_ and P_2_N_2_) were uninfluenced by the experimental tasks (all ps > 0.05).

The latency of the RotEPs (i.e., P_1_, mean 47 ±12ms) was not significantly different between the Updating and Control tasks (p = 0.64) or between the different rotation amplitudes (p = 0.60). The interaction Task x Amplitude was also not significant (p = 0.19).

### Dynamics of cortical source activity during spatial updating

The statistical maps computed over the 3 RotEP components (i.e., P_1_N_1_, N_1_P_2_ and P_2_N_2_) are shown in Figures 3-5. In these figures, warm color shadings indicate that the current computed in the Updating task was significantly greater than in the Control task. Cold color shadings indicate the opposite pattern. The MNI coordinates of maximal significant current difference between the Updating and Control tasks and their corresponding Brodmann areas are presented in Tables 1-3 for P_1_N_1_, N_1_P_2_ and P_2_N_2_, respectively.

**Table 1.** Maxima of regions showing significant differences between the Updating and Control conditions during the P_1_N_1_ interval of the RotEP (source space).

| Location | MNI coordinates |  |  | T-statistics | Brodmann area |
| --- | --- | --- | --- | --- | --- |
|  | X | Y | Z |  |  |
| left hemisphere |  |  |  |  |  |
| PPC | -35 | -60 | 51 | -3.09 | BA 39 |
| Lateral occipital | -50 | -76 | 17 | -2.33 | BA 19 |

**Table 2.** Maxima of regions showing significant differences between the Updating and Control conditions during the N_1_ P_2_ interval of the RotEP (source space).

| Location | MNI coordinates |  |  | T-statistics | Brodmann area |
| --- | --- | --- | --- | --- | --- |
|  | X | Y | Z |  |  |
| Left hemisphere |  |  |  |  |  |
| dIPFC | -15 | 48 | 47 | 3.47 | BA 8 |
| medial PFC | -13 | 47 | 10 | 2.77 | BA 10 |
| anterior medial PFC | -5 | 55 | -6 | 2.51 | BA 10 |
| SMA | -4 | 6 | 44 | 3.77 | BA 6 |
| medial temporal | -28 | -32 | -25 | 2.70 | BA 37 |
| Lateral occipital | -29 | -79 | 19 | -3.17 | BA 19 |
| Right hemisphere |  |  |  |  |  |
| dIPFC | 7 | 51 | 46 | 3.19 | BA9 |
| anterior medial PFC | 5 | 70 | 7 | 2.50 | BA10 |
| anterior PFC | 30 | 55 | 25 | 2.83 | BA 10 |
| SMA | 9 | 2 | 53 | 3.35 | BA 6 |
| temporal | 69 | -18 | -6 | 4.71 | BA 22 |
| medial temporal | 28 | -29 | -22 | 3.69 | BA 37 |

**Table 3.** Maxima of regions showing significant differences between the Updating and Control conditions during the P_2_N_2_ interval of the RotEP (source space).

| Location | MNI coordinates |  |  | T-statistics | Brodmann area |
| --- | --- | --- | --- | --- | --- |
|  | X | Y | Z |  |  |
| Left hemisphere |  |  |  |  |  |
| dIPFC | -7 | 52 | 44 | 2.49 | BA 9 |
| dIPFC | -2 | 24 | 51 | 3.24 | BA 8 |
| medial PFC | -2 | 50 | 45 | 2.44 | BA 9 |
| medial PFC | 3 | 17 | 50 | 3.03 | BA 8 |
| anterior medial PFC | -5 | 55 | -4 | 2.51 | BA 10 |
| SMA | -1 | 0 | 56 | 3.76 | BA 6 |
| medial temporal | -24 | -30 | -25 | 2.71 | BA 37 |
| temporal | -47 | 1 | -41 | 2.40 | BA 20 |
| PPC | -24 | -61 | 61 | 3.24 | BA 7 |
| precuneus | -1 | -78 | 46 | 2.6 | BA 7 |
| occipital | -1 | -94 | 19 | 4.83 | BA 18 |
| lateral occipital | -34 | -93 | 17 | -2.83 | BA 19 |
| Right hemisphere |  |  |  |  |  |
| dIPFC | 5 | 51 | 45 | 2.61 | BA 9 |
| dIPFC | 8 | 34 | 61 | 3.24 | BA 8 |
| medial PFC | 3 | 49 | 39 | 2.40 | BA 9 |
| medial PFC | 3 | 24 | 53 | 3.14 | BA 8 |
| SMA | 4 | -7 | 61 | 3.97 | BA 6 |
| premotor | 16 | -16 | 77 | 3.61 | BA 6 |
| medial temporal | 23 | -2 | -40 | 2.86 | BA 20 |
| PPC | 18 | -46 | 80 | 3.84 | BA 7 |
| Occipital | 2 | -87 | 19 | 4.65 | BA 18 |

**Figure 3.**
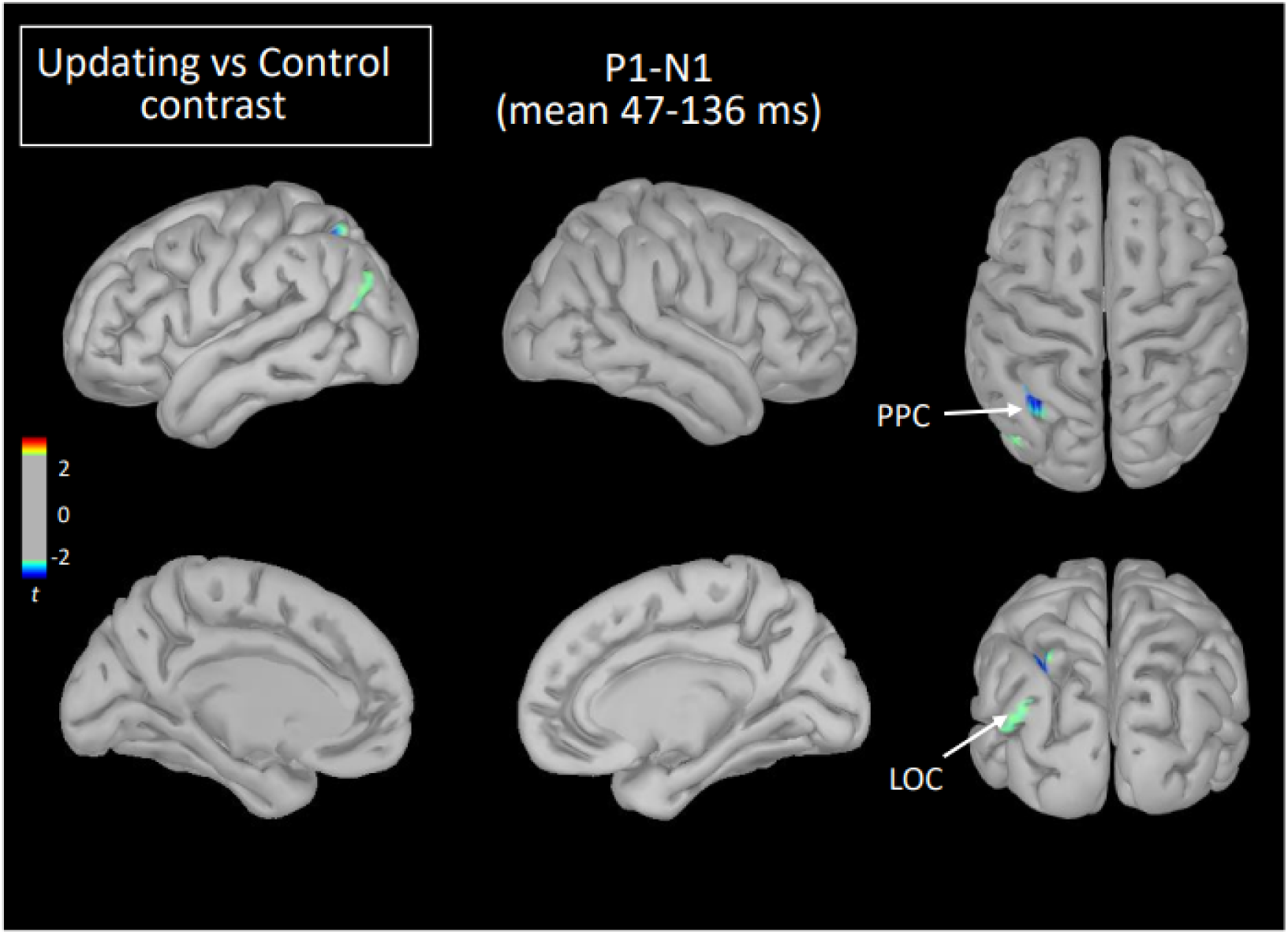
Statistical map (cortex only) for the Updating vs. Control contrast computed during the P_1_N_1_ component of the RotEP. LOC = lateral occipital cortex, PPC = posterior parietal cortex.

The mean current measured over the P_1_N_1_ interval, that is, between 47 ms and 136 ms after rotation onset, was strikingly alike between the Updating and Control tasks. The statistical map indicated that the cortical activity did not significantly differ between both tasks except for small areas of the left PPC and left lateral occipital cortex (LOC) (see Fig. 3). In these areas, the activity was smaller in the Updating than in the Control task.

Extensive differences in cortical current between both tasks emerged, however, in the second RotEP component (N_1_P_2_, between 130 ms and 303 ms after rotation onset). Overall, the cortical activity increased when participants tracked their initial position during the body rotations (see Fig 4). Significant task-related activities were mainly source-localized in the frontal and temporal cortices. Specifically, the supplementary motor complex (SMC), the dorsolateral prefrontal cortices (dlPFC), and the right anterior prefrontal cortex all showed greater activation in the Updating than in the Control tasks. Significantly greater current in the Updating task was also found in the right temporal cortex and in both medial temporal cortices, and in a small area of the right PPC. Only sparse regions of the LOC showed greater activation in the Control task than in the Updating task.

**Figure 4.**
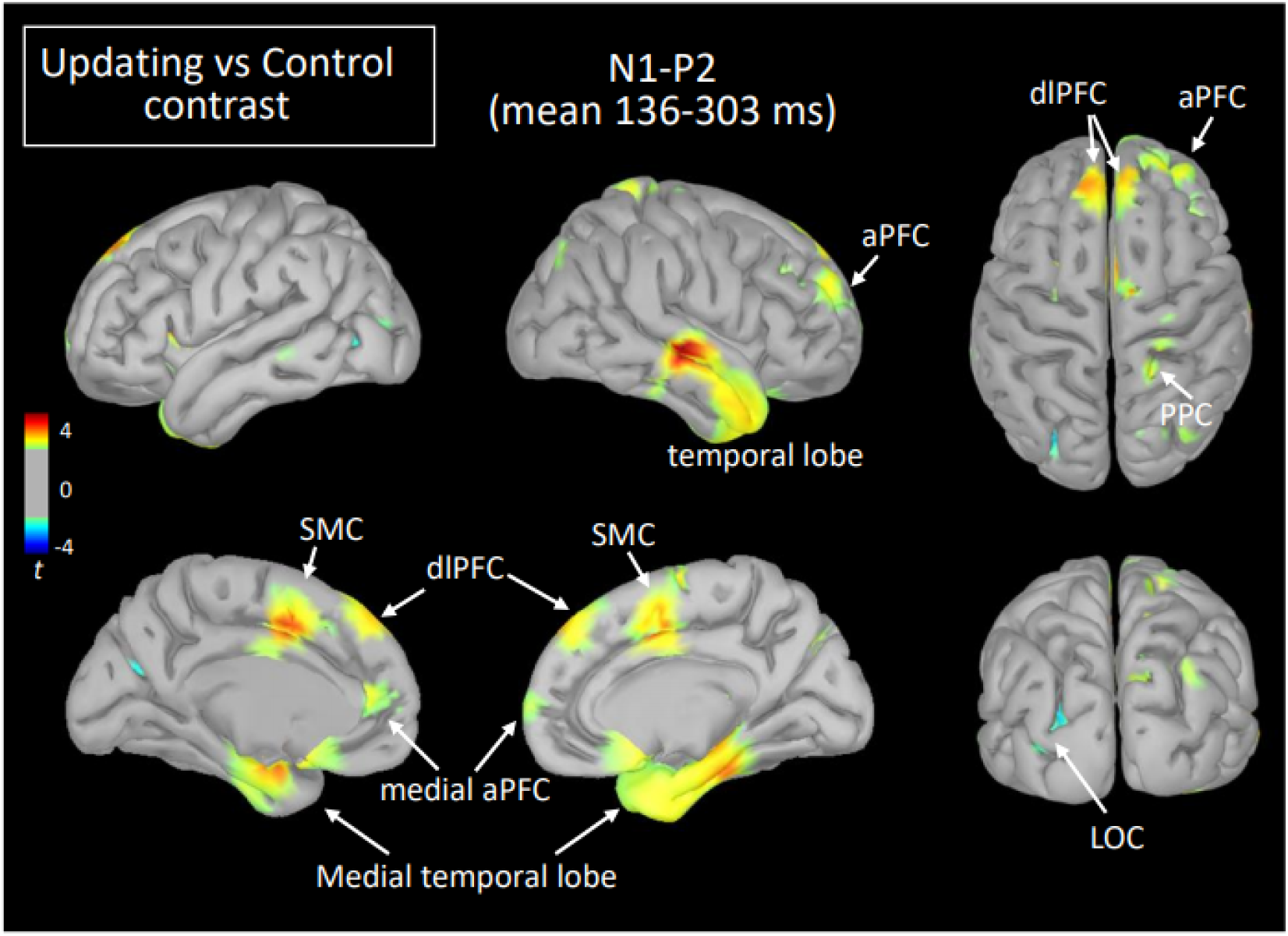
Statistical map (cortex only) for the Updating vs. Control contrast computed during the N_1_P_2_ component of the RotEP. aPFC — anterior prefrontal cortex, dIPFC = dorsolateral prefrontal cortex, LOC = lateral occipital cortex, PFC - prefrontal cortex, PPC = posterior parietal cortex, SMC = supplementary motor complex.

The increased activation observed in the Updating task during the N_1_P_2_ interval persisted in several cortical regions during the last RotEP component (P_2_N_2_, between 303 ms and 470 ms after rotation onset). This was the case for the bilateral dlPFC, SMC, medial temporal cortex, and for the right PPC (see Fig. 5). Other regions showed significant task-related activities exclusively in this last analysed interval of the RotEP. These regions were the right dorsal premotor areas, the left and right cuneus, the left precuneus, and the left PPC. Increased in the Updating task during N_1_P_2_, the activity of the right temporal lobe was no longer altered by the spatial updating processes during P_2_N_2_. The left LOC continued to exhibit greater activities in the Control task.

**Figure 5.**
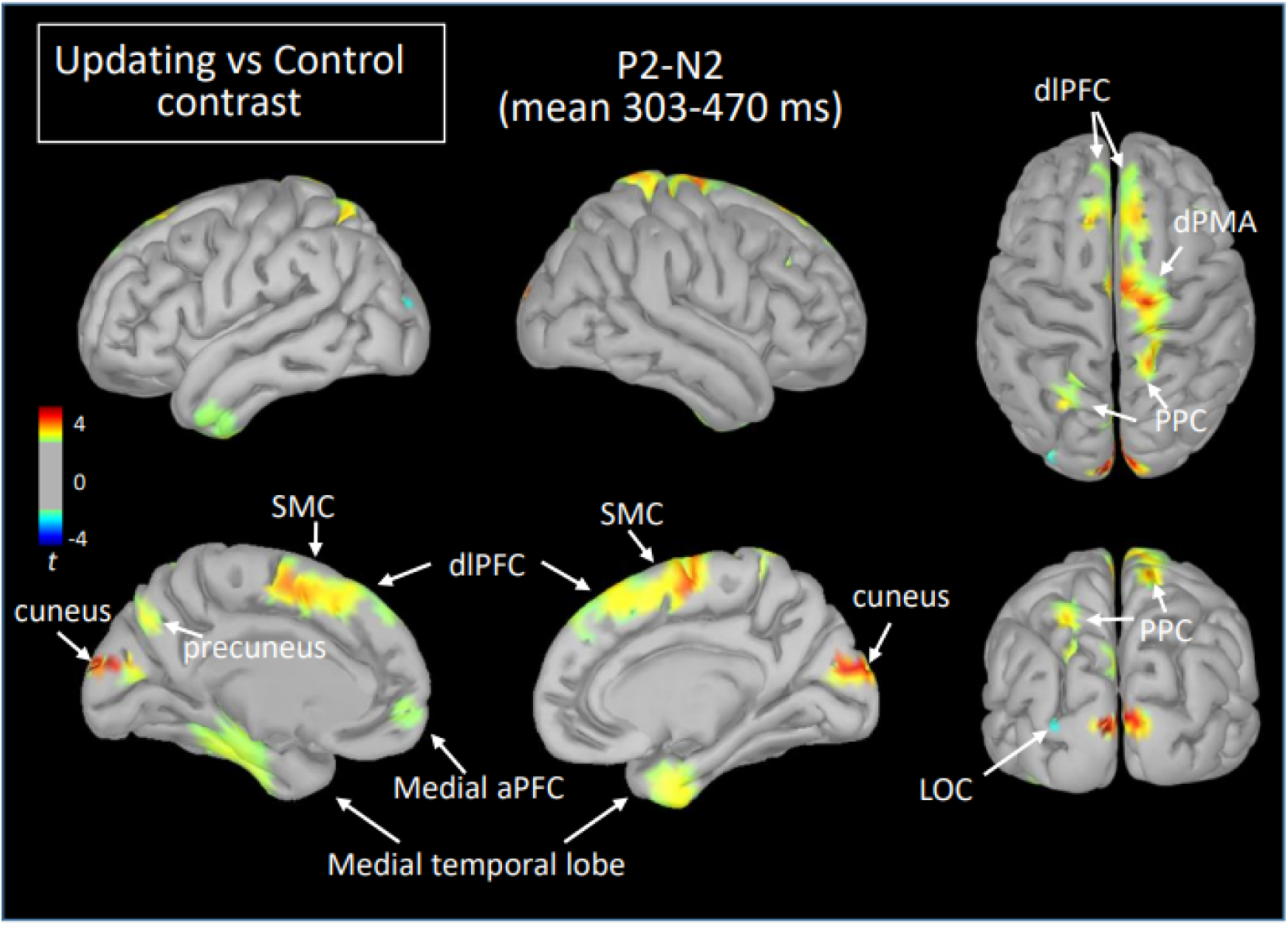
Statistical map (cortex only) for the Updating vs. Control contrast computed during the P_2_N_2_ component of the RotEP. aPFC = anterior prefrontal cortex, dlPFC = dorsolateral prefrontal cortex, dPMA = dorsal premotor area, LOC = lateral occipital cortex, PPC = posterior parietal cortex, SMC = supplementary motor cortex.

## Discussion

The present study was designed to gain insight into the dynamics of the cortical activations underpinning spatial updating during body motions. We used a protocol in which seated participants indicated their initial orientation after being passively rotated by different amplitudes in the dark. By contrasting, in successive time windows, the EEG activity recorded during body rotations from the EEG activity recorded in a baseline control task, we were able to isolate a discrete set of cortical areas involved in spatial updating processes and appraised their dynamics.

### The early processing of rotation-related cues is largely independent of the spatial updating

As a first salient finding, the current measured in the source space remained largely similar between the Updating and Control tasks over the first component of the RotEPs, a period spanning 47-136 ms after body rotation onset. The only significant effect revealed by the statistical map during this interval was a smaller activation in the Updating task in a few vertices of the left PPC and LOC. This effect persisted at both mid- and long latencies only for the LOC. The narrowness of the regions showing this trend raises the question of the robustness of this finding. Note, however, that the LOC contributes to the allocentric coding of space (Committeri et al., 2004; Zaehle et al., 2007, Ruotolo et al., 2019). Hence, the smaller activity observed in the LOC during the Updating task may have hampered the use of this frame of reference for encoding home position during body rotations. This could have indirectly enhanced the use of an egocentric frame of reference, which appeared more relevant in the present Updating task performed in the dark.

The absence of large task-related activation during P_1_N_1_ suggests that the first wave of idiothetic cue processing during the body rotations was not strictly linked to spatial updating. Yet, despite their non-specific nature, brain processes during the early phase of the rotations most certainly remained critical for spatial updating. This could be the case of the processes associated to vestibular inputs, which are the main carrier of body motion information during passive displacements in the dark (Valko et al., 2012).

The scarcity of early task-related activation was somewhat unexpected given that overt and covert attention influences neural activities related to the early processing of sensory cues (<100 ms; Di-Russo & Spinelli, 1999; Woldorff et al., 1987). One explanation for the lack of different cortical activations between the Updating and Control tasks may be that participants also directed their attention towards self-motion information in the latter task. This could have helped them to keep fixation on the chair-fixed LED during the rotations.

### Evidence of processes related to spatial working memory and spatial updating at mid-latencies

Large significant task-related activities emerged during the second RotEP component (i.e., N_1_P_2_, 136-303 ms after rotation onset). These activities were source-localized in a large network comprised mainly of the frontal, temporal and parietal regions. Like most task-related activities observed in the present study, the current measured in these regions were greater in the Updating than in the Control tasks.

As observed by Hartley et al. (2003) and Spiers & Maguire (2006) during wayfinding in virtual environments, the dlPFC and mPFC showed increased activations in the Updating task. The increased activations in these frontal areas could be associated to important executive functions for spatial processes. Their roles in maintaining spatial information in short-term working memory, and in the cognitive manipulation of information from the environment that is out of view (Wager & Smith, 2003; Gilbert & Burgess, 2008) could have been relevant in the present study for keeping track of the original orientation during body rotations.

On the other hand, the task-related activations observed in the aPFC and temporal cortices at both mid- and long latencies could be linked to the online processing of home direction during rotations. This interpretation is supported both by human studies showing that the aPFC is crucial to encoding spatial information about goals (Ekstrom et al., 2003; Ciaramelli, 2008; Spiers, 2008; see Poucet & Hok, 2017 for similar evidence in rodents) and by those reporting that the activities of the medial temporal lobe dynamically change according to the current distance of the spatial goal during navigation (Ekstrom et al., 2003; Kunz et al., 2021; Spiers & Barry, 2015). Together, the sustained activation observed in the prefrontal and temporal regions while participants were passively rotated could have then contributed to the maintenance of home orientation in working memory and in the encoding of its angular distance for use in the upcoming goal-directed pointing movements.

We also found significant increased activities in the Updating task in the right superior PPC during N_1_P_2_ which persisted bilaterally during P_2_N_2_. These results were to be expected given the well-recognized importance of the superior PPC for spatial processes, including those specifically linked to the updating of spatial representations during movements (Duhamel et al., 1992; Gutteling et al., 2015, 2016; Medendorp et al., 2003; Merriam et al., 2003; Pisella & Mattingley, 2004; Ventre-Dominey & Vallée, 2007). The increased activation observed in the superior PPC could also be related to the maintenance of spatial attention (see Ikkai & Curtis, 2011, for a review), which is a critical cognitive process for spatial updating. It should be noted that our observation that the increased activity in the right PPC preceded the increased activity over the left PPC is consistent with studies suggesting a right-hemispheric dominance for visual processes and remapping (Corbetta et al., 2000; Marshall et al., 2001; Pisella et al., 2011).

The SMC was the last region where significant task-dependent activities were observed at mid-latencies. These increased activities also lasted until the final analysis time window (i.e., P_2_N_2_). The SMC comprises the supplementary motor area (SMA), the supplementary eye fields (SEF) and the pre-supplementary motor area (preSMA) (Nachev et al., 2008). The SEF might have played a crucial role in the Updating task. This is suggested by studies showing that lesions affecting the SEF impair the accuracy of saccades towards a memorized visual target only if the patients are rotated before triggering the saccade (Pierrot-Deseilligny et al., 1993; 1995). The SEF was first considered as an oculomotor area (Schlag & Schlag-Rey 1987). Studies in Monkey, however, have identified a large population of SEF neurons that increase their activities during arm movements (40% of 337 neurons in Fujii et al., 2001; 42% of 106 neurons in Mushiake et al., 1996). These arm-related cells provide grounds for a plausible contribution of the SEF to providing relevant spatial information in the present study for pointing, after the rotations, towards the original body orientation.

### Long-latency activities during spatial updating could be linked to spatial representations

Bilateral increased activations were observed at long-latency (P_2_N_2_) in the cuneus, which is part of the medial visual cortex. In the present study, the only available visual input was a chair-fixed LED which the participants fixated throughout the trials. Because this visual input was present in both tasks, the contrast of Updating and Control tasks most likely cancelled out the neural activity evoked by the LED fixation. This possibility is well supported by the fact that the latency of the task-related activity observed in the cuneus (>300 ms) was much longer than the latency of visual-evoked potentials recorded in occipital lobe either by EEG or magnetoencephalography (<80 ms, Ellemberg et al., 2003; Lebar et al., 2015; Vianni et al., 2001). More likely, the late activity of the visual cortex may have resulted from non-visual top-down signals (mediated for instance by parietal or frontal regions, Michelli et al., 2004). This could have enabled the use of a visual-like representation to encode initial direction during the rotation, perhaps through visual mental imagery (see Kosslyn et al., 1999; Strokes et al., 2009). The fact that visual mental imagery activates the earliest visual cortex (BA 17 and 18) (Slotnick et al., 2005) affords this possibility.

It is interesting to note that the task-related activities observed in the medial surface of the left PPC (i.e. precuneus) at long latencies are consistent with the use of an egocentric frame of reference for encoding home position (Byrne et al., 2007; Chadwick et al., 2015; Wolbers et al., 2008). Based on idiothetic and gaze direction cues (Jeannerod, 1991; Paillard, 1987), this frame of reference appears most relevant in the present spatial task for updating body orientation during rotations in the dark. The egocentric spatial information contained in the precuneus would serve in contexts with body displacement but not for the mere egocentric judgments of objects location in steady body conditions (Chadwick et al., 2015). We noted that the increased activation of the precuneus observed in the Updating task was left lateralised. This lateralisation was also found by Chadwick et al. (2015) when using searchlight analysis to characterise neural activity in an fMRI navigation study, but not in their follow-up analyses using a more liberal threshold. Bilateral activation of the precuneus was also observed in the Wolbers et al.’s (2008) fMRI study when participants indicated the position of memorized objects following self-displacements in virtual environments. Note, however, that the low temporal resolution of fMRI scanners does not allow to determine whether activations in both hemispheres occur simultaneously. The results of the present EEG study may suggest that processes related to egocentric representations may have shorter latencies in the left than in the right precuneus, especially when encoding spatial positions from the contralateral visual hemifield.

Dorsal motor areas of the frontal lobe were strongly activated in the last analysed rotation interval of the Updating task. These areas have direct connections with spinal motor neurons (Chouinard & Paus, 2009). Importantly, however, we found that the increased activations in the dorsal motor areas were circumscribed to the right hemisphere, which was ipsilateral to the pointing arm. Although a fraction of ipsilateral connections reaches the spinal level (Kuypers, 1981), the absence of significant task-related activities in the contralateral dorsal motor areas suggests that the functions linked to their activations were more cognitive than motoric in nature. The marked activities found in the right dPMC in the Updating task is consistent with studies showing evidence of functional hemispheric differences between the left and right dPMC, with a salience for the right dPMC for spatial working memory processes (Jonides et al., 1993; Smith & Jonides, 1999). The dPMC could have beneficiated from inputs sent by prefrontal areas which showed task-related activities with shorter latencies (136-303 ms vs 303-470 ms after rotation onset). Indeed, the dPMC, and particularly the areas with only sparse projections to motoneurons, have dense connections with the PFC (Genon et al., 2017, Lu et al., 1994).

### Enhanced P_1_N_1_ amplitude during relevant idiothetic stimulation

The P_1_N_1_ component of the RotEP, as measured here over the vertex, had a significantly greater amplitude when the participants had to update their orientation during body rotations. This finding indicates that the amplitude of the cortical potential recorded at scalp level is a reliable marker of cognitive process enhancement related to spatial updating (including processes related to attention). This task-related effect on the P_1_N_1_ amplitude has found little echo, however, in the cortical current measured in the source space during the same P_1_N_1_ interval. This could be explained by the fact that the latter variable was obtained by averaging the cortical current over the P_1_N_1_ interval and that N_1_ marked the upper bound of this interval. The differential effect of the updating task on these two EEG measurements (i.e., P_1_N_1_ amplitude and current in the P_1_N_1_ interval) might suggest that the processes specifically dedicated to space updating essentially started with a latency close to that of N_1_, measured here as 136 ms after body rotation. Computing this latency with respect to the first cortical arrival of idiothetic cues (i.e., P_1_: 47 ms after rotation onset), probably provides a better estimate of the latency of the updating processes. Viewed from this perspective, our EEG recordings suggest that the cortical processes specifically involved in spatial updating have a latency of ∼89 ms.

### Limitations

Some limitations in the present study should be considered. We were careful to ensure that task-related cortical activities observed in the Updating task could not merely reflect motor-related processes linked to the preparation of the pointing movements produced after body rotations. With this in mind, we restrained our analyses on the early spatial updating processes, that is those with latencies <470 ms with respect to the onset of body rotation (i.e., N_2_ component of the RotEP). Cortical activities involved in spatial updating that had greater latencies could therefore not be identified. Also, because the present vestibular memory-contingent task involved goal-directed arm movements, task-related activities observed here could be, in part, specific to spatial updating in contexts of motor actions, and perhaps more specifically of goal-directed arm movements.

### Conclusion

The excellent temporal resolution of the EEG recordings combined with the enhanced spatial resolution of EEG data by sources analyses allowed us to obtain insight into the dynamics of spatial updating during body motions. Within the time window of our analyses (period spanning 47-470 ms after body rotation onset), virtually all task-related cortical changes of cortical activities during spatial updating involved increased rather than decreased activations. We found that the cortical activities specifically related to spatial updating during body rotation started ∼90 ms after the first arrival of idiothetic inputs to the cortex. The spatial updating processes were largely mediated by a fronto-temporo-posterior network and implicated more regions of the right hemisphere. Among the first cortical regions showing task-related activities were those that contribute to the encoding of spatial goals and to spatial working memory processes (e.g., aPFC, dlPFC, temporal cortices). The regions showing later task-related activities are known to be involved in the visual processing and in the egocentric representation of the environment (e.g., cuneus, precuneus). Because the spatial updating processes investigated here served as a basis for planning goal-directed arm movements, in future research, it will be interesting to contrast the present results with those obtained in tasks requiring other motor outputs (e.g., saccade, locomotion) or pure cognitive estimates of object locations during or after body displacements.

## Author contributions

**Jean Blouin:** Conceptualization, Methodology, Investigation, Data processing, Data curation, Writing-Original draft preparation, Reviewing, Validation. **Jean-Philippe Pialasse:** Conceptualization, Methodology, Software, Validation. **Laurence Mouchnino:** Conceptualization, Methodology, Reviewing, Validation. **Martin Simoneau**: Conceptualization, Methodology, Software, Investigation, Data processing, Writing, Reviewing, Validation.

